# Control of enhancer activation in the type I interferon response by the histone demethylase Kdm4d / JMJD2d

**DOI:** 10.1101/2023.01.14.524033

**Authors:** Rohit Chandwani, Terry C. Fang, Scott Dewell, Alexander Tarakhovsky

## Abstract

Transcriptional activation depends on the interplay of chromatin modifiers to establish a permissive epigenetic landscape. While histone 3 lysine 9 (H3K9) methylation has long been associated with gene repression, there is limited evidence to support a role for H3K9 demethylases in gene activation. Here we describe the H3K9 demethylase Kdm4d/JMJD2d as a positive regulator of type I interferon responses. In mouse embryonic fibroblasts (MEFs), depletion of JMJD2d attenuates the transcriptional response, conferring increased viral susceptibility, while overexpression of the demethylase results in more robust IFN activation. We find that the underlying mechanism of JMJD2d in type I interferon responses consists of an effect both on the transcription of enhancer RNAs (eRNAs) and on dynamic H3K9me^2^ at associated promoters. In support of these findings, we establish that JMJD2d is associated with enhancer regions throughout the genome prior to stimulation but is redistributed to inducible promoters in conjunction with transcriptional activation. Taken together, our data reveal JMJD2d as a chromatin modifier that connects enhancer transcription with promoter demethylation to modulate transcriptional responses.

## INTRODUCTION

Host defense is crucial to the survival of a species. In the type I interferon response, IFN-α/β (IFN) is secreted to initiate complex innate and adaptive defense against bacterial pathogens and viruses. At the heart of this response is the coordinated activation of a specific transcriptional program. Signaling networks downstream of the cytoplasmic pattern recognition receptors (PRRs) RIG-I and MDA-5 lead to the activation of IRF3/IRF7, NF-κB, and AP-1 that aggregate in the IFN-β enhanceosome.^1,2,3^ The coordinated activation of multiple transcription factors leads to the necessary and sufficient enhanceosome constituents for transcription of IFN-β.

Chromatin events are crucial to the elaboration of type I interferons. Transcription factor binding leads to subsequent docking of the GCN5 histone acetyltransferase complex and lysine acetylation at the *Ifnb1* promoter.^4^ Nucleosomal remodeling follows and results in engagement of the CBP-Pol II holoenzyme and transcription following recruitment of bromodomain-containing proteins (BRDs) and PTEFb.^5^

Recent work in our laboratory and others have implicated repressive lysine methylation as a key mediator of the magnitude of the type I interferon response. The presence of H3K9me^2^ at IFN and interferon-stimulated gene (ISG) loci correlates with a diminished response in fibroblasts as compared to cells of hematopoietic origin.^6^ In addition, GLP (a G9a histone methyltransferase homolog) represses ISG transcription by way of its enzymatic activity.^7^ These findings raise the question of whether there are corresponding histone demethylases that contribute to type I interferon signaling, specifically by facilitating the removal of repressive lysine methylation.

Here, we demonstrate that the histone 3 lysine 9 demethylase JMJD2d modulates the magnitude of type I interferon responses in mouse embryonic fibroblasts (MEFs) treated with exogenous nucleic acid and RNA viruses. We also show that JMJD2d largely associates with enhancers in unstimulated cells, but transcriptional activation of IFN-β and ISGs leads to redistribution from enhancers to inducible promoters. This shift in JMJD2d occupancy with stimulation is accompanied by H3K9 demethylation at inducible promoters and is lost with knockdown of JMJD2d. Together, these observations implicate JMJD2d in the regulation of the type I interferon response via an effect on the interface between the activity of distal regulatory elements and the removal of repressive promoter lysine methylation central to transcriptional activation.

## RESULTS

### JMJD2d is a positive regulator of the type I interferon response

To investigate a role for an H3K9 demethylase in the type I interferon response, we searched for H3K9 modifiers that were induced by dsRNA analog poly I:C. No specific acetyltransferase was inducible (data not shown). Of the demethylases tested, the mRNA transcript of the histone lysine demethylase JMJD2d was induced 10-fold by poly I:C (Fig. 1a) and to a lesser extent by other stimuli (LPS, IFN) (Fig. S1). To determine if JMJD2d has an effect on poly I:C-induced responses, we stably overexpressed JMJD2a and JMJD2d in MEFs and detected a significant increase in the expression of *Ifnb1* transcript with overexpression of JMJD2d but not JMJD2a in response to poly I:C (Fig. 1b). Conversely, JMJD2d depletion resulted in a ∼50% reduction in *Ifnb1* and *Ccl5* expression as compared to control siRNA-treated MEFs (Fig. 1c). To characterize the scope of the effect of JMJD2d depletion on the type I IFN response, we performed microarray analysis, identifying 113 genes induced by poly I:C in wild-type MEFs (Fig. 1d); several well-established interferon-stimulated genes (ISGs) were induced, including *Ifnb1, Mx1, Ccl5*, and the *Ifit* genes, and confirmed by RT-qPCR (Fig. S2). Broadly, ISGs displayed a significant attenuation of poly I:C-induced expression with JMJD2d-depletion in microarray analysis (Fig. 1e), with several targets confirmed by RT-qPCR (Fig. S3). While only ∼30% (34/113) of poly I:C stimulated genes were reduced at least 50% (Fig. 1f), nearly all genes were attenuated to some degree (Fig. 1g) suggesting a moderate but robust effect on the type I interferon response.

**Figure 1.**
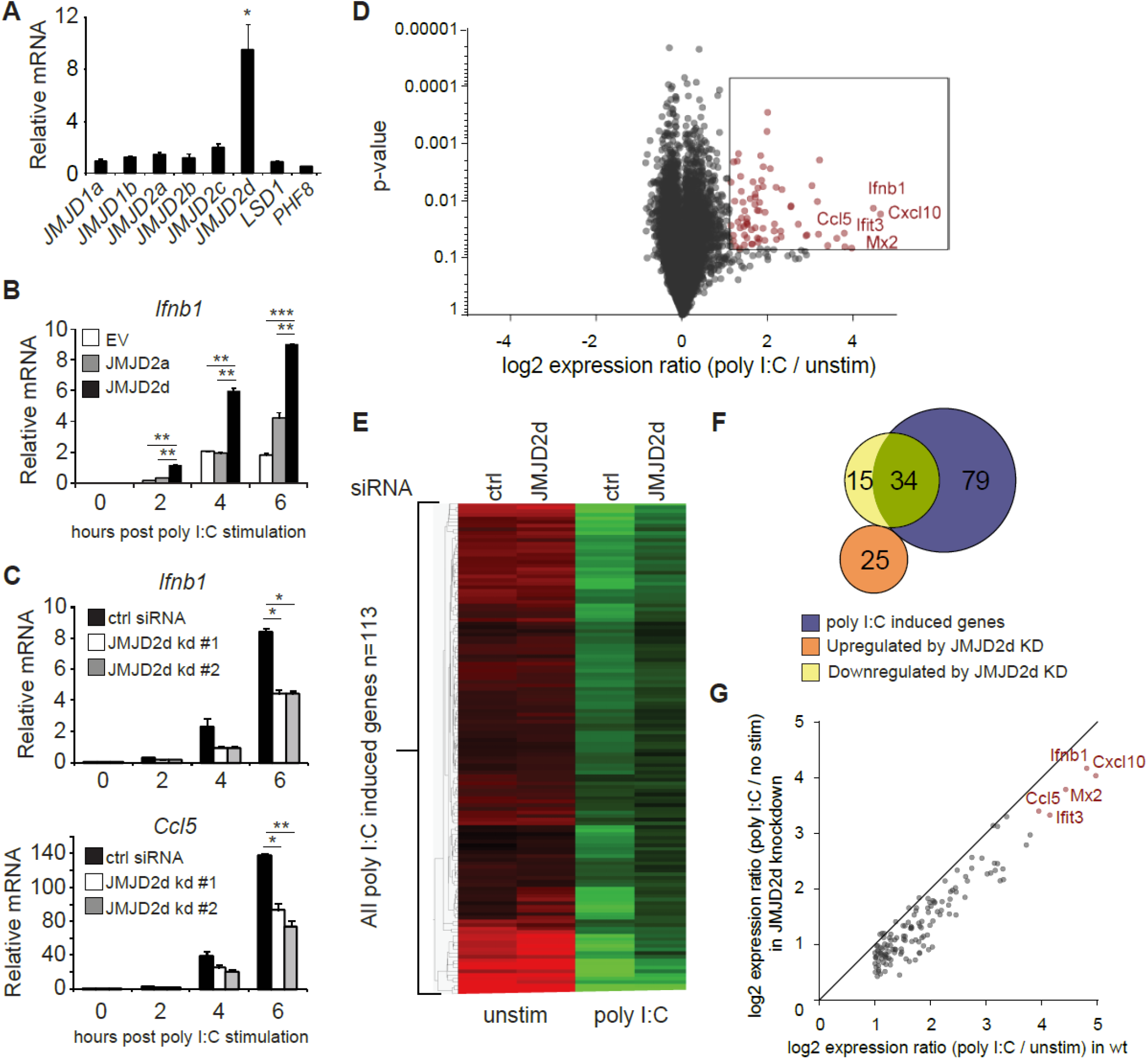
Modulation of the type I interferon response by the histone demethylase JMJD2d. (A) Relative expression of H3K9 demethylases in poly I:C-stimulated versus unstimulated MEFs. (B) Quantitative RT-PCR of *Ifnb1* induced by poly I:C in MEFs overexpressing empty vector (EV), JMJD2a, or JMJD2d. (C) Quantitative RT-PCR of *Ifnb1* and *Ccl5* induced by poly I:C in control or JMJD2d-depleted MEFs at the times indicated. (D) Volcano plot of poly I:C-inducible genes. Highlighted in red are 113 transcripts induced with log2 fold change > 1 and p<0.05. (E) Heatmap of the 113 poly I:C inducible genes in unstimulated (‘no stim’) and poly I:C transfected MEFs either treated with control (‘ctrl’) or JMJD2d (‘D2d’) siRNA. (F) Venn diagrams indicating overlap between poly I:C-inducible genes and transcripts either up-or down-regulated by depletion of JMJD2d. (G) Scatter plot indicating inducibility of 113 poly I:C inducible in the absence (x-axis) or presence (y-axis) of JMJD2d knockdown.

### JMJD2d modulates the antiviral response

Next, we evaluated the effect of JMJD2d modulation in the context of viral infection. In response to vesicular stomatitis virus (VSV) infection, we found that JMJD2d-depleted MEFs displayed attenuated upregulation of the *Ifnb1* transcript (Fig. 2a), while JMJD2d-overexpressing MEFs expressed 2-fold more *Ifnb1* mRNA (Fig. 2b). Remarkably, we found the small effect on *Ifnb1* to have a substantially larger effect on the frequency of viral infection, as knockdown of JMJD2d increased viral susceptibility to VSV-GFP approximately 2.5-fold, while overexpression resulted in a 2.5-fold increase in resistance to a Sindbis-mCherry virus (Fig. 2c).

**Figure 2.**
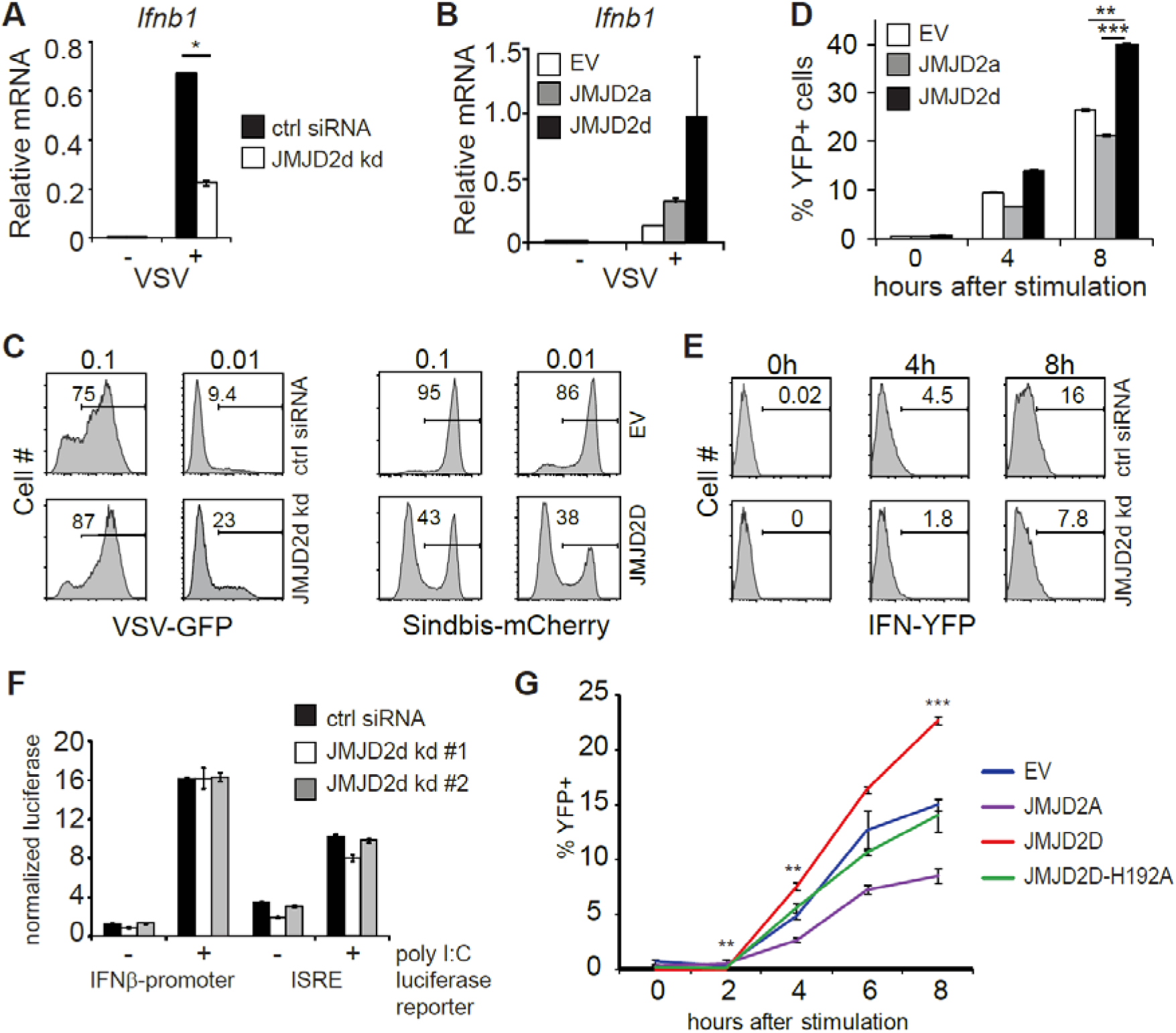
JMJD2d controls the stochastic response to virus. (A,B) Quantitative RT-PCR of *Ifnb1* induced by VSV after 12h infection in (A) control or JMJD2d-depleted MEFs and in (B) MEFs overexpressing empty vector (EV), JMJD2a, or JMJD2d. (C) Histograms indicating percentages of GFP^+^ or mCherry^+^ cells following infection with VSV-GFP and Sindbis-mCherry virus, respectively, at the indicated MOIs in either control or JMJD2d-depleted MEFs. (D) Percentages of YFP^+^ cells following stimulation with poly I:C at the indicated times in either EV, JMJD2a, or JMJD2d-overexpressing IFN-YFP MEFs. (E) Histograms indicating percentages of YFP^+^ cells following stimulation with poly I:C at the indicated times in either control or JMJD2d-depleted IFN-YFP MEFs. (F) Normalized luciferase expression in MEFs treated with control or JMJD2d siRNA, transfected with either *Ifnb1*-luc or ISRE-luc reporters, and stimulated with poly I:C. (G) Percentages of YFP^+^ cells following stimulation with poly I:C at the indicated times in either EV, hJMJD2A, or hJMJD2D or hJMJD2D-H192A overexpressing IFN-YFP MEFs. *p<0.05; **p<0.01; ***p<0.001.

The elaboration of IFN-β displays stochastic features in many cell types, such that only a small percentage of cells will upregulate *Ifnb1* transcripts and subsequent protein.^8^ We therefore asked if JMJD2d affects the frequency of cells producing *Ifnb1*. To this end, we utilized MEFs derived from reporter mice in which the yellow fluorescent protein (YFP) is expressed from a bicistronic mRNA linked by an internal ribosomal entry site (IRES) to the endogenous *Ifnb1* mRNA.^9^ Recapitulating our findings for total transcript, we observed that overexpression of JMJD2d increased the frequency of IFN-β cells by 50%, whereas overexpression of JMJD2a had no such effect (Fig. 2d). Conversely, JMJD2d depletion by siRNA decreased the percentage of IFN-β+ MEFs induced by poly I:C by ∼50% (Fig. 2e). We also observed a similar phenomenon in the context of Sendai virus infection with substantial abrogation in the percentage of YFP+ cells following infection (Fig. S4).

### The effect of JMJD2d on IFN and ISG activation requires chromatin and enzymatic function

To interrogate the mechanisms required for this effect of JMJD2d on the type I interferon response, we asked if the presence of chromatin was necessary. To exclude a signaling defect in JMJD2d-depleted cells, we generated luciferase reporter constructs conjugated to the *Ifnb1* promoter (IFN-luc) or to an IFN-sensitive response element (ISRE-luc) that were transfected into MEFs, providing non-chromatinized substrates. We observed that siRNA-mediated knockdown of JMJD2d did not attenuate poly I:C-induced activation of either reporter (Fig. 2f), suggesting that the effects of JMJD2d depletion are not due to an impact on signaling leading to transcription factor engagement. To assess the requirement for the enzymatic function of JMJD2d, we mutated the catalytic histidine residue in the JmjC domain, generating the H192A mutant. As before, IFN-YFP MEFs engineered to overexpress wild-type JMJD2D doubled in the frequency of YFP-positivity after poly I:C, while those expressing the catalytic dead JMJD2D were similar to the empty vector (EV) alone (Fig. 2g), suggesting that demethylase activity is required for our observed phenotype.

### JMJD2d predominately associates with enhancers in the genome

Given the requirement of chromatin and demethylase activity, we hypothesized that the effects of JMJD2d modulation on type I IFN signaling require histone 3 lysine 9 demethylation. To address this question directly, we evaluated the genome-wide occupancy of JMJD2d. Using JMJD2d-3xFLAG MEFs, we were able to identify a significant enrichment in JMJD2d occupancy (as compared to empty vector [EV] control) at enhancer positions flanking the *Ifnb1* and *Ccl5* loci that was not present at promoters (Fig. 3a). Of note, the binding pattern of JMJD2d was relatively broad, occurring over multiple kilobases. Genome-wide integration of ChIP-seq data showed substantial enrichment of JMJD2d (FLAG) signal above EV at enhancers and not promoters (Fig. 3b). Using algorithms optimized for the broad-type enrichment that we observed (see Methods), we identified all JMJD2d-binding sites in the genome. With a false-discovery rate set to <2.5%, there were 4821 discrete JMJD2d-binding sites throughout the genome in unstimulated JMJD2d overexpressing MEFs; of these binding sites, approximately 70% colocalized with enhancer positions, while only 8% overlapped with promoters (Fig. 3c; left pie chart). Of the top 1000 JMJD2d binding sites in the genome, 85% colocalized with enhancer positions (Fig. 3c; right pie chart), suggesting that JMJD2d is strongly associated with enhancers.

**Figure 3.**
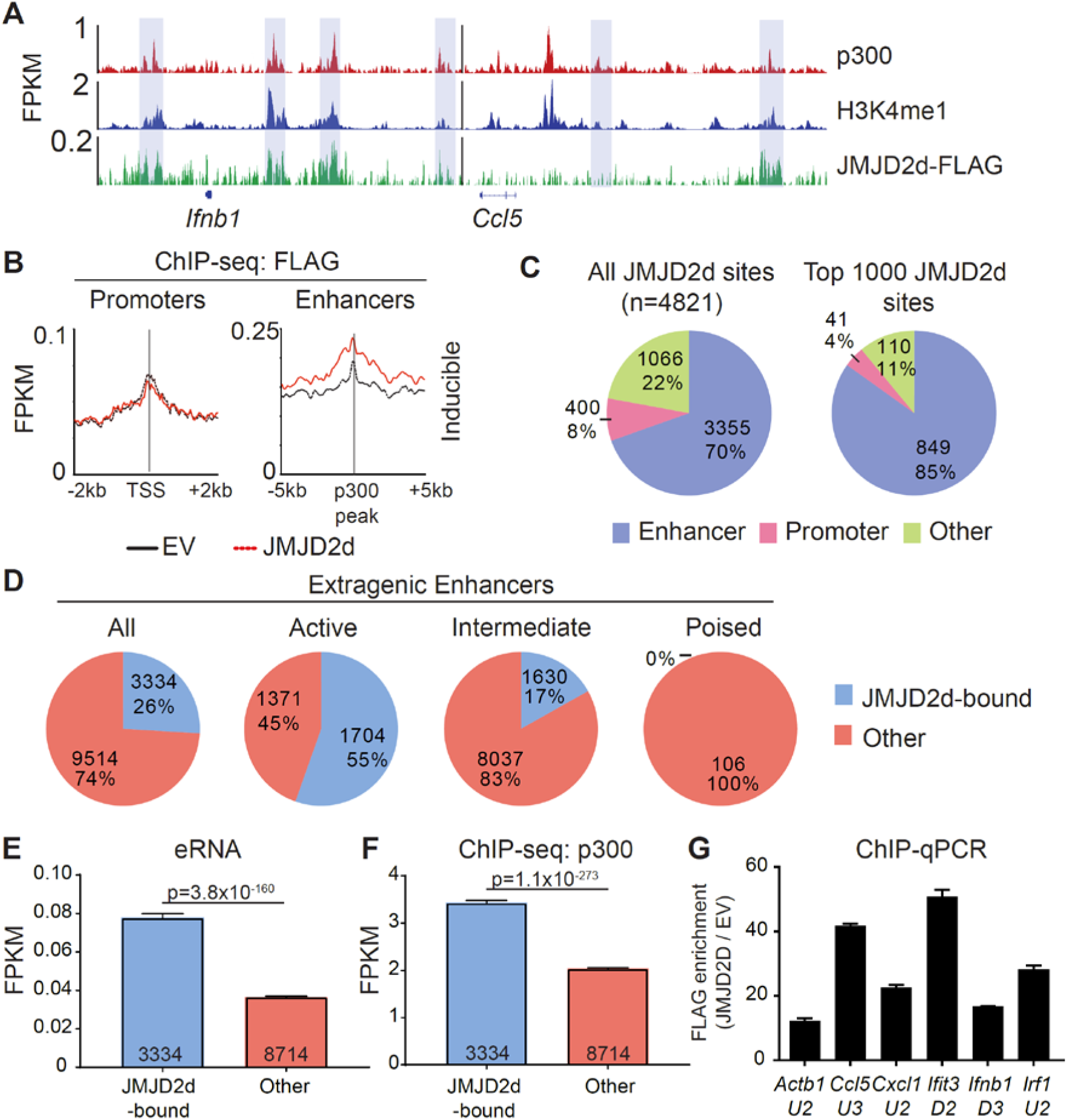
JMJD2d colocalizes with active enhancers in the genome. (A) ChIP-sequencing profiles for p300, H3K4me^1^, and Flag-tagged JMJD2d in unstimulated MEFs at the *Ifnb1* and *Ccl5* loci. (B) Integrated profile plots of ChIP-seq data representing aggregate enrichment of JMJD2d at poly I:C-inducible promoters or enhancer-promoter unit (EPU)-assigned inducible enhancers in unstimulated MEFs. (C) Percentage of either all JMJD2d sites or top 1000 JMJD2d binding sites in the genome that colocalized with enhancer, promoter, or other regions, in unstimulated MEFs. (D) Percentage of active (H3K9ac), intermediate (H3K9-), and poised (H3K9me^3^) enhancers that colocalize with JMJD2d binding sites. (E,F) Average FPKM from (E) whole-transcriptome RNA-seq data and (F) p300 ChIP-seq data for JMJD2d-bound and other enhancers. (G) ChIP-qPCR indicating enrichment of JMJD2D at the shown enhancer locations.

We next asked if JMJD2d is specifically enriched at particular types of enhancers, as we suspected that JMJD2d might be bound to H3K9me^2/3^+ enhancers given that this is the substrate for lysine demethylation. Curiously, however, JMJD2d was much more frequently associated with H3K9-acetylated rather than H3K9-methylated regulatory elements (Fig. S5). Using the typical classification for enhancer activity^10^, we found that JMJD2d was much more frequently found associated with active (H3K27ac+) rather than poised (H3K27me^3^+) enhancers (Fig. 3d). Among all extragenic enhancers, JMJD2d-bound elements were noteworthy for higher levels of enhancer RNA (Fig. 3e) and p300 enrichment (Fig. 3f) suggesting that JMJD2d occupancy correlates strongly with enhancer activity, a finding we confirmed in a chromatin state hidden Markov model (Fig. S6). Notably, several of the most active enhancers surrounding type I IFN response genes were particularly enriched for JMJD2d as compared to an enhancer found in proximity to the *Actb* locus (Fig. 3g). Together, these findings suggest that JMJD2d associates with active enhancers in steady-state MEFs.

### Chromatin dynamics in the type I interferon response

Next, we extensively profiled the range of dynamic epigenetic modifications in response to poly I:C by ChIP-seq, hypothesizing that we would find stimulus-induced demethylation of H3K9me^2/3^ at enhancers. At the *Ifnb1* and *Ccl5* loci, we found poly I:C-induced accumulation of H3K4me^3^, H3K9ac, H4ac, and Pol II at promoters and transcriptional start sites (TSSs) and H3K36me^3^ in gene bodies (Fig. 4a). Robust induction of these marks and increased transcription was seen across the 113 inducible genes (Fig. 4b-c), but not a panel of 113 randomly selected genes (Fig. S7). We also noted the presence of several regulatory elements [blue rectangles] in the vicinity of these genes enriched in the characteristic enhancer features

**Figure 4.**
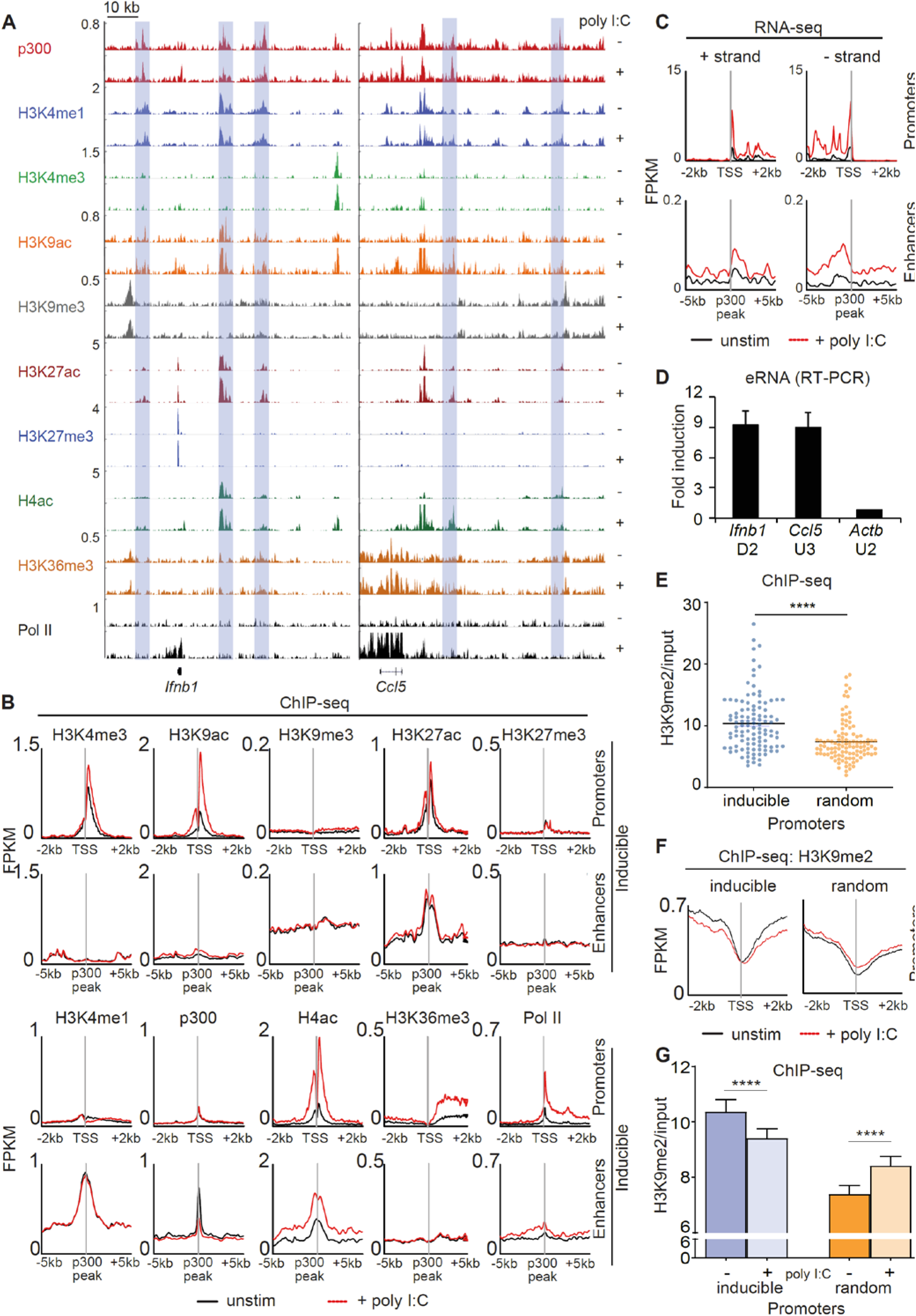
Dynamic activation of enhancers occurs in the type I interferon response. (A) ChIP-sequencing profiles corresponding to the antibodies as listed in unstimulated (-) and poly I:C-transfected (+) MEFs at the *Ifnb1* and *Ccl5* loci. Blue rectangles denote enhancers (H3K4me1 and p300 enrichment in the absence of H3K4me3). (B+C) Integrated profile plots of ChIP-sequencing data (B) or strand-specific RNA-sequencing data (C) representing aggregate enrichment in either poly I:C-inducible genes (‘promoters’) or their EPU-designated associated enhancers (‘enhancers’) before and after treatment with poly I:C. (D) Relative expression of eRNAs in poly I:C-stimulated versus unstimulated MEFs. (E) Vertical scatter plot of H3K9me2 abundance (normalized to input) at the promoters of 113 poly I:C-inducible (‘inducible’) or 113 randomly selected (‘random’) genes. (F) Integrated profile plots of ChIP-sequencing data for H3K9me2 at poly I:C-inducible or random genes, before and after treatment with poly I:C. (G) Bar chart quantifying H3K9me2 (normalized to input) at the promoters of inducible and random genes before and after stimulation with poly I:C. ****p<0.0001

H3K4me^1^ and p300 (Fig. 4a).^10,11,12^ Extragenic enhancers were annotated genome-wide and assigned to our set of 113 inducible genes based on the enhancer-promoter unit (EPU) method, which incorporates proximity and the location of nearby insulator (CTCF) binding sites (see Supplemental Methods).^13^ Broadly, we evaluated the enhancers assigned to all 113 poly I:C-inducible genes (see Methods), observing stimulus-induced accumulation of active histone marks -namely, H4ac and Pol II, but not p300, H3K4me1, or H3K27ac (Fig. 4b). Importantly, extragenic enhancers belonging to the aforementioned random set of genes did not show increases in active histone marks (Fig. S8). At specific enhancers, we also found H3K9me^3^ uniquely occupied ‘flanking’ positions at the enhancer that ranged in distance from 3 to 6 kb from the p300 peak^14^, but we were not able to detect loss of this H3K9me^3^ with stimulation.

To further characterize our annotated set of inducible enhancers, we examined enhancer RNA arising from these regulatory elements. eRNA has been described as transient species originating from active enhancers (H3K27ac^+^, H3K9ac^+^) at the ‘center’ (p300 peak) with bidirectional transcription and variable polyadenylation.^15-16^ Multiple lines of evidence suggest that eRNAs act as a platform for the recruitment of additional molecules such as Mediator and cohesin central to enhancer-promoter communication by ‘looping’ or ‘tracking’ mechanisms.^17-19^ Using strand-specific RNA-sequencing, enhancer RNA within EPUs of poly I:C-inducible genes were ostensibly induced, whereas enhancers in EPUs of random genes were not similarly activated (Fig. 4c; Fig. S9). Using qRT-PCR for eRNA, we were able to detect ∼10-fold induction of specific eRNAs upstream and downstream of the *Ifnb1* and *Ccl5* gene but not at eRNAs of uninduced genes (Fig. 4d).

Next, we asked whether promoter H3K9me^2^ is a dynamic repressive mark in the type I interferon response, as we had described previously in fibroblasts.^6^ Consistent with our prior report, we were able to find that poly I:C-inducible genes in MEFs were indeed marked by high levels of H3K9me^2^ at promoters (relative to random genes) (Fig. 4e). H3K9me^2^ at promoters was also reduced with poly I:C stimulation in the inducible set but not the random gene set (Fig. 4f), a finding that was statistically significant (Fig. 4g). These findings thus indicated that demethylation of promoter-associated H3K9me^2^ is a feature of the type I interferon response.

### JMJD2d is associated with inducible promoters upon stimulus-induced transcription

Given that promoter-associated H3K9me^2^ is lost from inducible genes, we hypothesized that JMJD2d might be responsible for demethylation at these locations. However, in unstimulated fibroblasts, JMJD2d predominately colocalized with enhancers. To reconcile these findings, we interrogated the dynamics of JMJD2d binding in the type I interferon response to determine if JMJD2d can be implicated in promoter events. Following the stimulation of MEFs with poly I:C, we noted increased JMJD2d signal throughout the coding sequence of the *Ifnb1* and *Ccl5* genes (Fig. 5a, yellow rectangles). More importantly, promoters throughout the genome globally showed enrichment of JMJD2d (above EV control) after stimulation that was not present prior (Fig. 5b). As before, all JMJD2d binding sites were identified throughout the genome, and a much greater percentage overlapped with promoters following stimulation than prior (13% vs 8% for all sites; 12% vs 4% for top 1000 sites) (Fig. 5c). More precisely, JMJD2d occupancy was observed at ∼40% of inducible genes, while random genes were not frequently associated with JMJD2d (Fig. 5d). We observed that the specific JMJD2d signal from the five most highly induced genes (*Ifnb1, Cxcl10, Mx2, Ifit3, Ccl5*) was especially increased (Fig. 5e) and that this stimulation-induced rise was robust (∼4-fold) across all inducible genes to the exclusion of random genes (where a decrease in JMJD2d occupancy was detected) (Fig. 5f). These data show that JMJD2d is recruited to the promoters of inducible genes in the type I interferon response.

**Figure 5.**
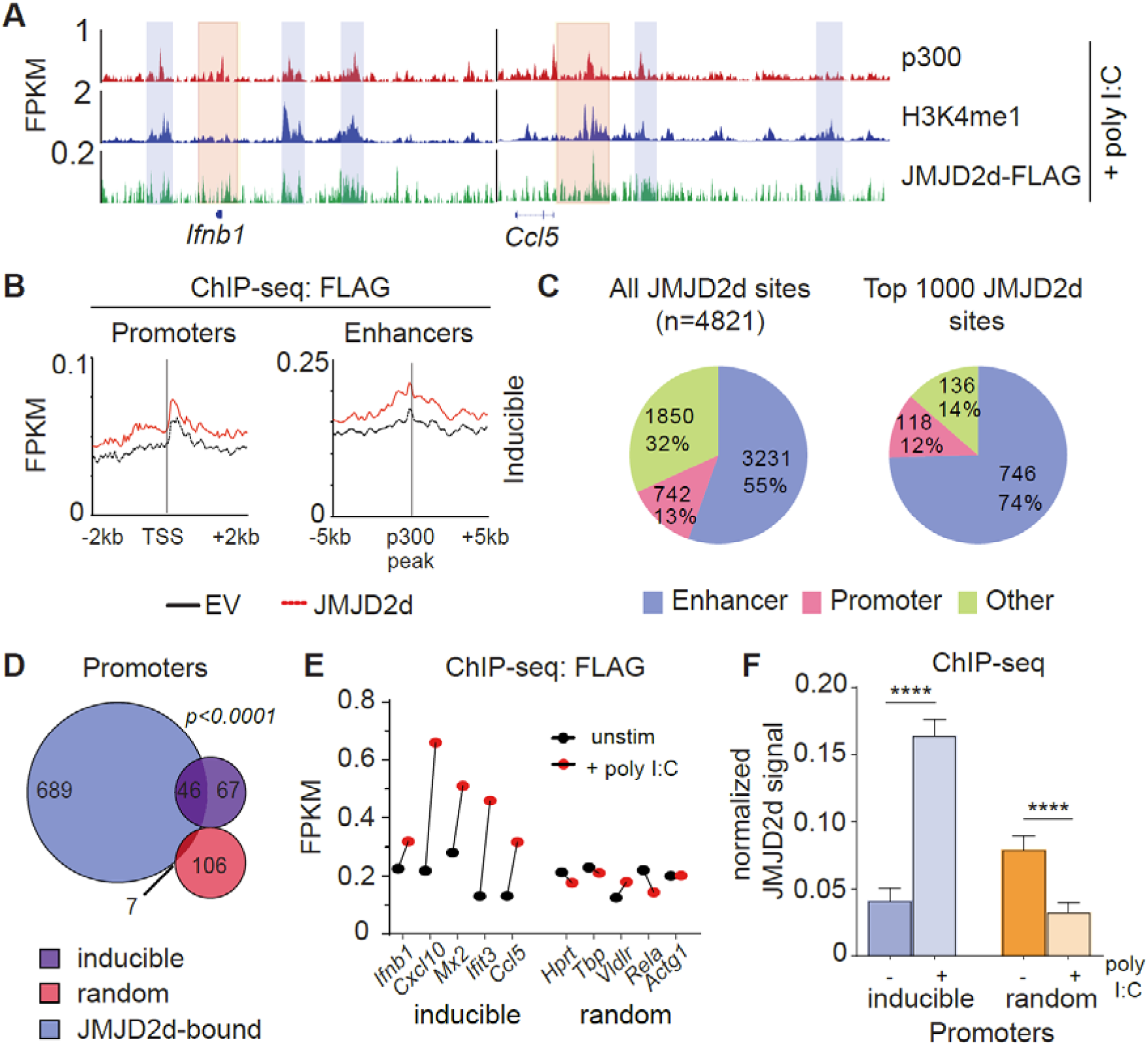
JMJD2d accumulates at inducible promoters in the type I interferon response. (A) ChIP-sequencing profiles for p300, H3K4me^1^, and Flag-tagged JMJD2d in poly I:C-transfected MEFs at the *Ifnb1* and *Ccl5* loci. (B) Integrated profile plots of ChIP-seq data representing aggregate enrichment of JMJD2d at poly I:C-inducible promoters or enhancer-promoter unit (EPU)-assigned inducible enhancers in poly I:C-treated MEFs. (C) Percentage of either all JMJD2d sites or top 1000 JMJD2d binding sites in the genome that colocalized with enhancer, promoter, or other regions, in poly I:C-stimulated MEFs. (D) Venn diagrams indicating overlap between JMJD2d-bound promoters in poly I:C stimulated MEFs and poly I:C inducible or random genes. Fisher’s exact test used to compare proportions between inducible and random sets. (E) FPKM of ChIP-seq signal for FLAG in JMJD2d-3xFLAG MEFs at the indicated promoters prior to (black circles) or after poly I:C-treatment (red circles). (F) Bar charts of normalized JMJD2d signal in all inducible or random promoters prior to or after treatment with poly I:C. ****p<0.0001

### JMJD2d depletion abrogates eRNA transcription and promoter demethylation

Given the association of JMJD2d with regulatory elements, we next sought to ascertain if JMJD2d depletion was associated with any specific changes to enhancer chromatin. To this end, we profiled chromatin marks in poly I:C stimulated MEFs with and without JMJD2d depletion, focusing on the extragenic enhancers annotated to inducible genes. In this group of genomic locations, we saw that enhancers of inducible genes did not change in their occupancy of p300 or the amount of H3K4me^1^, H3K9ac, or H3K9me^3^ (Fig. 6a). By contrast, we did observe a modest reduction in Pol II and H4Ac enrichment at the enhancers of inducible but not random genes (Fig. 6b), suggesting an impact on eRNA transcription. In support of these findings, we confirmed that JMJD2d knockdown attenuates the expression of particular eRNA species associated with key type I interferon response gene loci (*Ifnb1, Ccl5*, and *Mx2*) (Fig. 6c). These findings thus point to a role for JMJD2d in the activation of enhancers and in the transcription of eRNA.

**Figure 6.**
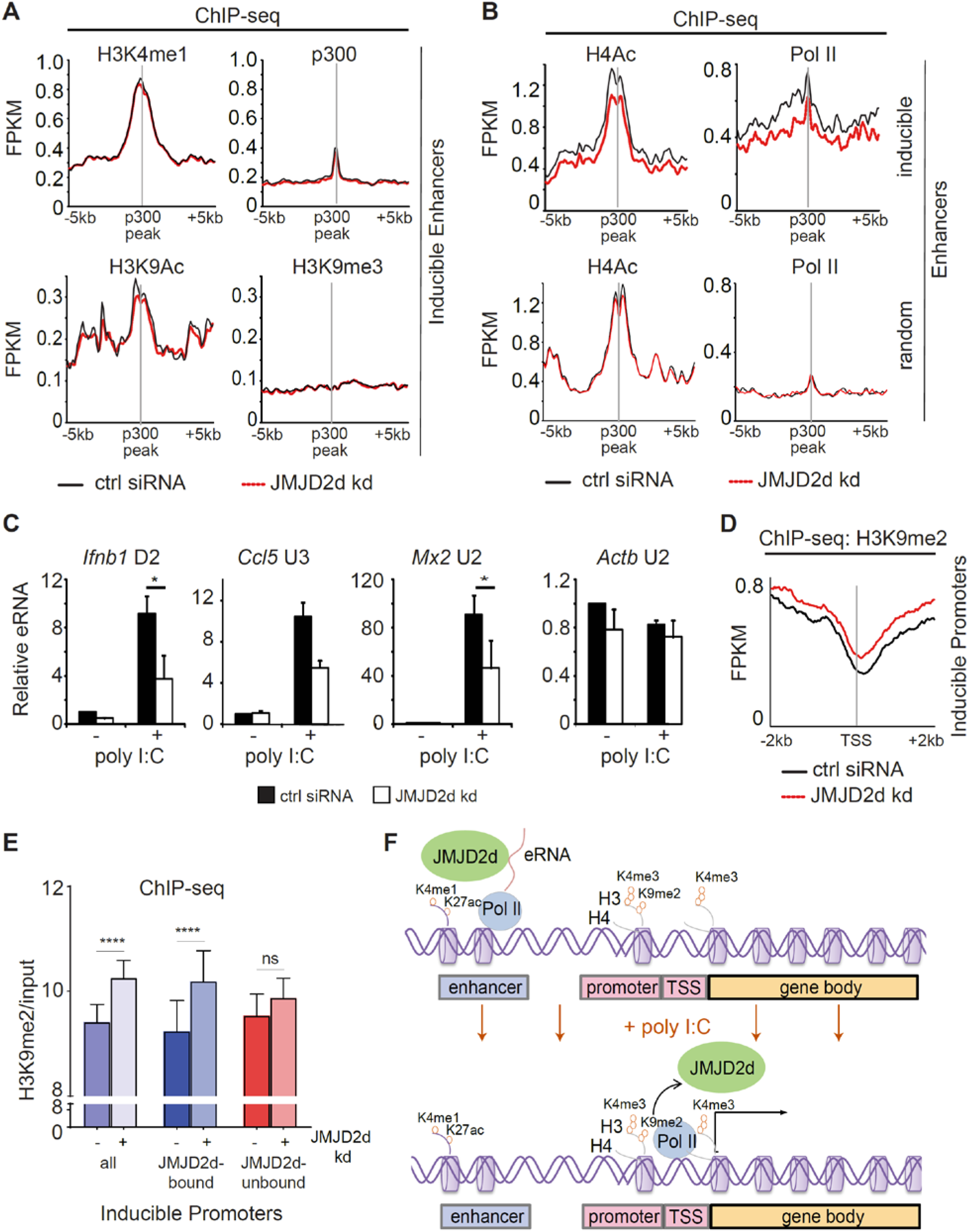
Enhancer transcription and promoter methylation are modulated by JMJD2d. (A,B) Integrated profile plots of ChIP-sequencing data representing aggregate enrichment of the indicated marks at enhancers of poly I:C-inducible genes, and (B) at enhancers of both poly I:C inducible and random genes, in cells treated with poly I:C in the absence and presence of JMJD2d knockdown. (C) Relative expression of eRNAs in unstimulated or poly I:C-treated control or JMJD2d-depleted MEFs. (D) Integrated profile plot of ChIP-sequencing data indicating aggregate enrichment of H3K9me2 at inducible promoters in poly I:C stimulated MEFs in the presence or absence of JMJD2d knockdown. (E) Bar chart quantifying H3K9me2 (normalized to input) at the promoters of inducible genes, or subsetted on the basis of JMJD2d-binding, in poly I:C stimulated cells +/-JMJD2d knockdown. (F) Schematic depicting JMJD2d at active (H3K4me1^+^ H3K27ac^+^) and transcribed (eRNA^+^) enhancers and at inducible (H3K4me3^+^ Pol II^+^) promoters after stimulation, where H3K9me2 is reduced.

Finally, we interrogated whether JMJD2d loss had any implications for dynamic H3K9 methylation observed at promoters of inducible genes. Via ChIP-seq, we observed that JMJD2d depletion is associated with increased H3K9me^2^ levels at inducible promoters in the context of a poly I:C-induced response (Fig. 6d). To determine if this was specific to the presence of JMJD2d, we separated JMJD2d-bound (n=46) from JMJD2d-unbound (n=67) promoters (Fig. 5d). Quantification of H3K9me^2^ levels showed that promoter methylation is increased in the context of JMJD2d depletion primarily at those promoters normally bound by the demethylase (Fig. 6e). These findings thus serve to implicate JMJD2d as an enhancer-bound histone 3 lysine 9 demethylase that controls eRNA transcription and is delivered to promoters where it demethylates H3K9me^2^ to facilitate stimulus-induced transcription (Fig. 6f).

## DISCUSSION

In summary, our observations show that enhancer chromatin is markedly dynamic in the innate immune response with inducible accumulation of active histone marks and eRNA expression. We find that the H3K9 demethylase JMJD2d tightly associates with enhancers in the genome and modulates the transcription of eRNA and target gene expression in response to poly I:C.

Our findings complement prior work identifying H3K9me^3^ as a histone mark that controls enhancer activity in the inflammatory response.^20^ JMJD2d was suggested in this work to be the enzyme that modulates this enhancer-associated H3K9me^3^, but here we find no such large-scale dynamics of H3K9me^3^. This is largely attributable to our unbiased approach to enhancer analysis; we consider all poly I:C-inducible gene-associated enhancers rather than only those high in JMJD2d or associated with select genes. We suspect that in our conditions there exists a subset of JMJD2d-enriched enhancers associated with specific genes that have altered H3K9me^3^ – certainly, these could include the *Ifnb1, Ccl5*, and *Mx2* enhancers, as these were most strongly affected by JMJD2d.

Importantly, we find that the vast majority of chromatin-bound JMJD2d specifically associates with enhancers suggesting that it is a specific regulator of enhancer chromatin. As a whole, however, our analysis shows that only a minority of all enhancers (30.5%) are bound by JMJD2d. This points to a role for JMJD2d in the regulation of only a subset of enhancers, perhaps to confer cell-type specificity; in turn, this implies that enhancers not bound by JMJD2d have restricted function in certain cell types. Moreover, we find that JMJD2d binds most active enhancers and accumulates at the most highly upregulated enhancers, suggesting the enzyme is potentiate maximal gene activation by maintaining lysine demethylation. How JMJD2d is recruited to specific enhancers and its possible interactors on chromatin (including transcription factors, coregulators, and Mediator and cohesin complexes) remain to be elucidated, but these findings newly implicate a demethylase in the complex events between enhancer activation and gene transcription.^22-23^

Lastly, our data identify a role for JMJD2d in the control of the innate immune response. Most notably, JMJD2d depletion affected only ∼30% of poly I:C-inducible genes, suggesting JMJD2d has the potential to attenuate but not abrogate transcriptional responses, providing specificity as a therapeutic target. JMJD2d may function differently depending on cell type because enhancer chromatin is more varied across cell types than is the promoter landscape^12,21^, leading to differential JMJD2d targeting. To the extent that we have also demonstrated JMJD2d occupancy is dynamic with poly I:C stimulation, we suspect that the signaling context is also likely to alter the effect that JMJD2d targeting may have. We conclude that JMJD2d is a regulator of type I interferon responses and that its function on enhancer chromatin implies a role across a multitude of cellular contexts.

## ACKNOWLEDGMENTS

This work was supported by a National Institutes of Health KL2 Career Development Award (to RC).

## AUTHOR CONTRIBUTIONS

RC designed and performed experiments, analyzed data, and wrote the paper. TCF performed experiments and analyzed data. SD performed data analysis. AT supervised experiments, analyzed data, and supervised data analysis.

## METHODS

### Cell lines and culture

MEFs were generated from d13.5 embryos from C57/B6 and IFN-YFP mice by trypsinization of the embryonic body and culture of adherent cells. Immortalization was performed with SV40 latent T antigen. All MEFs were cultured in DMEM with 15% fetal bovine serum, 1% L-Glutamine, 1% penicillin-streptomycin, 1% non-essential amino acids, and 0.1% 2-mercaptoethanol (GIBCO). MEFs from passage number 12-15 were used for experimentation.

### Quantitative reverse-transcribed PCR

RNA was isolated with the RNeasy mini kit (Qiagen) or by the High Pure RNA isolation kit (Roche) and subsequently quantified using the NanoDrop 8000 spectrophotometer (Agilent). RNA was reversed transcribed using the Transcriptor First Strand Kit (Roche). qPCR was carried out using SYBR Green on a LightCycler 480 (Roche). Gene expression was displayed relative to HPRT.

### ChIP-sequencing

Briefly, cells were incubated as above and at the indicated times following stimulation or viral infection were treated with 1% formaldehyde for 10 minutes. Fixation was terminated with the addition of glycine to a final concentration of 0.125M. Cells were then washed serially with PBS/0.5% FCS and collected. Cells were then lysed to isolate chromatin and sonicated for 10-20 min (depending on cell type) using a Bioruptor (Diagenode). Fragmentation of sonicated chromatin to lengths of 200-500 bp was confirmed by agarose gel electrophoresis.

For immunoprecipitation, beads were prepared with 7-10μg of antibody coupled to M-280 mouse or rabbit Dynabeads (Invitrogen) for 8 hours as per the manufacturer’s instructions. Beads were then washed with PBS/0.5% FBS on a magnet and 20-50μg sonicated chromatin was added and incubated overnight on a rotator. Following immunoprecipitation, beads were washed in modified RIPA wash buffer (50 mM HEPES-KOH pH 7.6, 100 or 300 mM LiCl, 1 mM EDTA pH 8.0, 1% NP-40, 0.7% Na-Deoxycholate) and then in TE wash buffer (10 mM Tris-HCl pH 8.0, 1 mM EDTA pH 8.0, 50 mM NaCl). DNA was then eluted at 65°C for 40 min in elution buffer (50 mM Tris-HCl pH 8.0, 10 mM EDTA pH 8.0, 1% SDS). Crosslinks were reversed overnight and protein and RNA were digested. For validation of ChIP-seq, subsequent purified DNA was analyzed via qPCR.

For ChIP-seq library preparation, blunt-end DNA was prepared using the End-It End Repair Kit (Epicentre), “A” bases were added using Klenow fragment (NEB), and adapters for sequencing (Illumina) were ligated using T4 DNA ligase (NEB). Adapter-ligated DNA was then amplified using PE primers 1.0 and 2.0 (Illumina) and Phusion polymerase (ThermoScientific). Libraries were analyzed via agarose gel electrophoresis, a NanoDrop spectrophotometer, on an Agilent Bioanalyzer for integrity, appropriate size, and the presence of adapter dimers or primer dimers. Libraries were then loaded onto the Illumina HiSeq 2000 according to Illumina protocols.

### RNA-sequencing

RNA was isolated in TriZOL reagent (Invitrogen) and purified.. Purified RNA was quantified on a NanoDrop spectrophotometer (Agilent). Total RNA was depleted of small RNAs (miRNA, tRNA) using the RNeasy Mini Kit (Qiagen). Ribosomal RNA was depleted from 10μg total RNA using the Ribo-Zero Magnetic Kit (Epicentre). Ribo-depleted RNA was fragmented; cDNA synthesis and strand-specific library generation was carried out using the Script-Seq V2 Kit (Epicentre) and the FailSafe PCR enzyme mix. Libraries were loaded onto the Illumina HiSeq 2000.

### Cell transfections and stimulations

All knockdown experiments were performed using the Lipofectamine RNAiMAX transfection reagent (Invitrogen) with siRNA delivered at a concentration of either 20 or 50 nM in serum-depleted medium (OptiMEM; Invitrogen) as per the manufacturer’s instructions. Reverse transfections were employed with cells added to preformed siRNA:lipofectamine complexes pre-incubated for 20 minutes. Subsequent cell stimulations or viral infections were performed 48 hours after initial siRNA knockdown. Reporter constructs were transfected using the Lipofectamine 2000 transfection reagent (Invitrogen) according to the manufacturer’s instructions.

MEFs were stimulated by transfection of 2 µg/ml poly(I:C) into the cells using Lipofectamine 2000 transfection reagent (Invitrogen) following the manufacturer’s instructions. Alternatively, MEFs were stimulated with 500 U/ml recombinant IFN-β (R&D Systems) for the indicated times. To block IFN-β-induced signaling, MEFs were pre-incubated with 10 µg/ml IFNαR1 antibody (MAR1-5A3, eBioscience, catalogue number 16-5945).

### Retroviral transduction

Retrovirus was generated by transfecting BHK cells (ATCC) or Phoenix cells (laboratory stocks) with 1-4 μg plasmid using the Lipofectamine 2000 Transfection Reagent (Invitrogen). Supernatants were collected at 24 and 48 hours post-transfection and added to MEFs pre-cultured in six-well plates. Polybrene (Sigma) was added at a concentration of 8 μg/mL. Six well plates were then centrifuged at RT for 90 min and incubated at 37°C overnight; efficiency of transduction was measured by flow cytometry 48 hours following spin infection.

### Viruses and viral infections

Viruses were obtained as follows. Stocks of GFP-expressing VSV (designated VSV-GFP M51R), mCherry-expressing Sindbis virus, as well as influenza A (Puerto Rico/8/34 (H1N1)), amplified in 8-day old embryonated chicken eggs and titered by plaque assay on Madin-Darby canine kidney cells) were kindly provided by Adolfo Garcia-Sastre. Wild type VSV Indiana serotype (San Juan), originally a gift from Milton Schlesinger, was grown and titered on BHK-21 cells as previously described (Bick et al., 2003). Sendai virus was kindly provided by Charles Rice.

MEFs were infected with individual viruses with a multiplicity of infection (MOI) between 0 and 1 by diluting the virus stock in PBS containing 0.5% to 1% FBS. Infection was allowed to proceed for 1 h. After removal of the virus containing supernatant, cells were incubated in fresh medium. For detection of infected cells via FACS, cells were infected with a reporter virus that leads to the expression of GFP in the infected cells (VSV-GFP, sindbis-mCherry). Cells were analyzed for GFP or mCherry expression at the indicated times.

Plasmids and cloning. Plasmids expressing hJMJD2A and hJMJD2D were generated by cloning PCR-amplified constructs containing the endogenous Jmjd2a and Jmjd2d loci conjugated to 3xFLAG into the MigR1 plasmid backbone. All plasmids were transformed into competent E.coli at 42°C and subjected to antibiotic selection. Overnight bacterial cultures were grown, plasmids purified via Maxiprep (Qiagen), and quantified on a NanoDrop spectrophotometer (Agilent).

### Flow cytometry and cell sorting

For flow cytometry, cells were collected and fixed in PBS containing 1% paraformaldehyde and analyzed on a FACSCalibur flow cytometer (BD) using CellQuest software. For cell sorting, cells were collected and fixed in the same fashion but analyzed on a FACSAria (BD) cell sorter to divide into YFP+ and YFP-populations.

### Primers

JMJD2d, Ccl5, Mx2, IRF7, Ifit1, and Ifit3 TaqMan probes obtained from Applied Biosystems. All other primers were designed and obtained from Sigma and are summarized in Supplemental Tables 2 and 3. All forward and reverse primers were mixed to a final concentration of 50 μM.

### Microarray analysis

1-5 μg of total RNA from 2-3 MEF samples per group was used to prepare biotinylated RNA using the Ambion Illumina TotalPrep RNA Amplification Kit (Applied Biosystems) according to the manufacturer’s instructions. This RNA was hybridized to Illumina MouseRef-8 v2.0 expression BeadChip kits, with chips then scanned using the Illumina BeadArray Reader followed by analysis using Genespring (Affymetrix). The raw expression data was quality assessed and subjected to background adjustment and quantile normalization. The individual gene expression levels were compared by using an unpaired Student’s T-test (P<0.05) and by pairwise comparison.

### Antibodies

The following antibodies were employed: histone H3 antibody (Abcam, ab1791), Histone H3 (di methyl K9) antibody (Abcam, ab1220), ChIPAb+ Trimethyl-Histone H3 (Lys4) (Millipore, 17-614), Anti-acetyl-Histone H4 antibody (Millipore, 06-866), RNA polymerase II CTD repeat YSPTSPS antibody [4H8] -ChIP Grade (Abcam, ab5408), anti-histone H3 monomethyl K4 antibody (Abcam, ab8895), anti-histone H3 acetyl K9 antibody (Abcam, ab4441), anti-histone H3 trimethyl K9 antibody (Abcam, ab8898), anti-trimethyl-histone 3 (Lys27) (Millipore, 07-449), anti-histone H3 (acetyl K27) antibody (Abcam, ab4729), and anti-Flag M2 antibody (Sigma, F1804).

### Analysis of sequencing data

For all alignments, reads were aligned to the mm9 build of the mouse genome, downloaded from the UCSC Genome Browser. For ChIP-seq data, reads were aligned at 36bp (H3K4me3 and RNAPII) or 51bp (all others), allowing for only unique alignments with 2 mismatches using Bowtie v0.12.7. Sequencing reads from replicate ChIP experiments were combined for final analyses. For RNA-seq data, reads were 101bp and were aligned using Tophat v2.0. Reads were segmented to 25bp and 2 mismatches were allowed. Junctions were supplied from RefSeq annotations. Peak calling was performed as follows: MACS v1.4 was used for peak calling for p300 and CCAT3.0 for all others. Suitable inputs or controls were used for all analyses. Scores 60 and higher were used for MACS and FDR values of 0.05 or less were used for CCAT3.0. For profiling of ChIP-seq and RNA-seq data, reads were extended 100bp from the 3’ end to account for the expected size of the fragments and binned into 100bp windows. Windows upstream and downstream of indicated features were queried for 5kb, or the distances indicated. In libraries where the rate of duplication was high, duplicates were removed to avoid biasing profiles. Profiles are reported as reads per million mapped reads per bin/window size (100bp). Fragments-per-kilobase-per-million-mapped-reads values were calculated from alignment data for the intervals/peaks regions specified. A pseudocount of 1 read per 100M aligned reads per kilobase was used to avoid division-by-zero errors.

### Enhancer definition

To qualify as an enhancer, p300 peaks (with score 60 from MACS) were assessed based on the following criteria. A p300 peak could not intersect with a H3K4me3 peak called for any condition (L0/L4/2D0/2D4), could not be within 1kb of a RefSeq gene start site (TSS) or 2kb from the start site (TSS) of any spliced EST entries downloaded from the UCSC genome browser, and must intersect with a H3K4me1 peak location from any of the conditions. Intragenic enhancers were defined as those p300 peaks that passed these criteria that were contained within the transcriptional unit for RefSeq genes. Extragenic enhancers were those remaining enhancers that were not within 5kb of the transcription end site, to avoid strong RNAPII from biasing profiling signal. For eRNA profiling, additional filtering was performed to eliminate potentially un-annotated genes, repetitive sequence or other problematic features. As the characteristic profiles of eRNAs are short, less than 2kb transcripts in the region 2kb downstream on the plus strand and 2kb upstream on the minus strand were considered. The following were excluded: any transcript for which the ratio of reads contained on the forward strand 2kb downstream of the p300 peak summit to reads on the minus strand 2kb upstream was more than 5; any transcript for which the signal in the region +2-4kb or -2-4kb was higher than 2kb immediately upstream or downstream (to eliminate potential long/genic transcripts); any high signal over 1 FPKM that constituted a small fraction of loci with an aberrantly high number of alignments potentially due to repetitive signal; any loci with homology to any ribosomal RNA sequence. Inducible enhancers were defined as follows: Enhancers were deemed to be induced if H4ac or H3K9ac peaks were found in the poly I:C stimulated condition using the respective unstimulated H4ac or H3K9ac library as a control, using CCAT3.0 with an FDR < 0.1. These peaks had to intersect enhancers that passed all aforementioned criteria.

### Chromatin state analysis

ChromHMM was used to determine chromatin states.^24,25^ 29 states were specified, with the indicated target samples being used. Proper control libraries were used. The software models potential states using a hidden Markov model to determine likely combinations of enriched regions. Enriched regions for each target are then assessed for the representation in the various potential chromatin states. Target regions, such as genes, enhancers and other genomic loci can then be queried to ascertain the predominant chromatin states present at those loci.

## REFERENCES

1. Stetson DB, Medzhitov R. (2006) Type I interferons in host defense. Immunity 25: 373–381.

2. Thanos D, Maniatis T. (1995) Virus induction of human IFN beta gene expression requires the assembly of an enhanceosome. Cell 83: 1091–1100.

3. Panne D, Maniatis T, Harrison SC. (2007) An atomic model of the interferon-beta enhanceosome. Cell 129: 1111–1123.

4. Agalioti T, et al. (2000) Ordered recruitment of chromatin modifying and general transcription factors to the IFN-beta promoter. Cell 103: 667–678.

5. Jang MK, et al. (2005) The bromodomain protein Brd4 is a positive regulatory component of P-TEFb and stimulates RNA polymerase II-dependent transcription. Mol Cell 19: 523–534.

6. Fang TC, et al. (2012) Histone H3 lysine 9 di-methylation as an epigenetic signature of the interferon response. J Exp Med 209: 661–669.

7. Ea CK, Hao S, Yeo KS, Baltimore D. (2012) EHMT1 protein bingds to nuclear factor-κB p50 and represses gene expression. J Biol Chem 37: 31207–17.

8. Zhao M, Zhang J, Phatnani H, Scheu S, Maniatis T. (2012) Stochastic expression of the interferon-beta gene. PLoS Biol 10, e1001249.

9. Scheu S, Dresing P, Locksley RM. (2008) Visualization of IFNbeta production by plasmacytoid versus conventional dendritic cells under specific stimulation conditions in vivo. Proc Natl Acad Sci 105: 20416–20421.

10. Rada-Iglesias A, et al. (2011) A unique chromatin signature uncovers early developmental enhancers in humans. Nature 470: 279–283.

11. Ghisletti S, et al. (2010) Identification and characterization of enhancers controlling the inflammatory gene expression program in macrophages. Immunity 32: 317–328.

12. Heintzman ND, et al. (2009) Histone modifications at human enhancers reflect global cell-type-specific gene expression. Nature 459: 108–112.

13. Shen Y, et al. (2012) A map of the cis-regulatory sequences in the mouse genome. Nature 488: 116–120.

14. Zentner GE, Tesar PJ, Scacheri PC. (2011) Epigenetic signatures distinguish multiple classes of enhancers with distinct cellular functions. Genome Res 21: 1273–1283.

15. Kim TK, et al. (2010) Widespread transcription at neuronal activity-regulated enhancers. Nature 465: 182–187.

16. De Santa F, et al. (2010) A large fraction of extragenic RNA Pol II transcription sites overlap enhancers. PLoS Biol 8: e1000384.

17. Kagey MH, et al. (2010) Mediator and cohesin connect gene expression and chromatin architecture. Nature 46: 430–435.

18. Ren X, Siegel R, Kim U, Roeder RG. (2011) Direct interactions of OCA-B and TFII-I regulate immunoglobulin heavy-chain gene transcription by facilitating enhancer-promoter communication. Mol Cell 42: 342–345.

19. Engel N, Raval AK, Thorvaldsen JL, Bartolomei SM. (2008) Three-dimensional conformation at the H19/Igf2 locus supports a model of enhancer tracking. Hum Mol Genet 19: 3021–3029.

20. Zhu Y, van Essen D, Saccani S. (2012) Cell-type-specific control of enhancer activity by H3K9 trimethylation. Mol Cell 46: 408–423.

21. Visel A, et al. (2009) ChIP-seq accurately predicts tissue-specific activity of enhancers. Nature 457: 854–858.

22. Calo E, Wysocka J. (2013) Modification of enhancer chromatin: what, how, and why? Mol Cell 49: 825–837.

23. Kaikkonen MU, et al. (2013) Remodeling of the enhancer landscape during macrophage activation is coupled to enhancer transcription. Mol Cell 51(3):310–25.

24. Ernst J, et al. (2011) Mapping and analysis of chromatin state dynamics in nine human cell types. Nature 473: 43–49.

25. Kharchenko PV, et al. (2011) Comprehensive analysis of the chromatin landscape in Drosophila melanogaster. Nature 471: 480–485..

