## Supplemental Figures & Figure Legends for "Control of enhancer activation in the type I interferon response by the histone demethylase Kdm4d / JMJD2d"

**SUPPLEMENTAL MATERIAL**

**Supplemental Figure 1. Induction of JMJD2d in the type I interferon response.** Relative expression of JMJD2d in MEFs treated with the indicated stimuli, 4 hours following treatment.

**Supplemental Figure 2. Induction of IFN and ISGs by poly I:C in MEFs.** Normalized fold induction of *Ifnb1, Mx1, Tnf, Il6, Irf7, Ccl5, Mx3, Ifit1* and *Ifit3* in wild-type MEFs stimulated with poly I:C at the indicated times. All RT-qPCR data shows are normalized to expression of HPRT.

**Supplemental Figure 3. Depletion of JMJD2d diminishes the poly I:C-inducible expression of several ISGs.** Normalized expression of *Mx1*, *Mx2, Tnf, Il6,* and *Jmjd2d* following 48h treatment with the indicated siRNA and stimulation with poly I:C for the indicated times.

**Supplemental Figure 4. Depletion of JMJD2d decreases the percentage of IFN-producing cells in response to infection with Sendai virus.** Histograms indicating percentages of $YFP^+$ cells following infection with Sendai virus for the indicated times in either control or JMJD2d-depleted IFN-YFP MEFs.

**Supplemental Figure 5. JMJD2d associated with $H3K9ac^+$ enhancers.** Pie charts of JMJD2d binding frequency among H3K9ac, H3K9-, and $H3K9me^3$ enhancers.

**Supplemental Figure 6. Chromatin state analysis shows JMJD2d colocalized with active enhancers in chromatin.** Heatmap of hromatin state analysis with multivariate hidden Markov model emission showing 29 chromatin states based on ChIP-seq for the indicated marks. Intensity of each box indicates the frequency with which each state is accompanied by a histone mark.

**Supplemental Figure 7. Dynamic histone marks in the response to poly I:C is restricted to inducible but not random promoters.** Integrated profile plots of ChIP-sequencing data representing aggregate enrichment in either 113 poly I:C-inducible genes ('inducible') or a set of 113 randomly chosen genes ('random').

**Supplemental Figure 8. Dynamic histone marks in the response to poly I:C is restricted to enhancers associated with inducible but not random genes.** Integrated profile plots of ChIP-sequencing data representing aggregate enrichment in either extragenic enhancers within the EPUs of poly I:C-inducible genes ('inducible') or a set of randomly chosen genes ('random').

**Supplemental Figure 9. Dynamic eRNA transcription in the response to poly I:C is restricted to enhancers associated with inducible but not random genes.** Integrated profile plots of RNA-sequencing data representing expression of either extragenic enhancers within the EPUs of poly I:C-inducible genes ('inducible') or a set of randomly chosen genes ('random').

**Supplemental Table 1. List of poly I:C inducible genes.** Also given are indicated fold changes 4 hours after treatment with poly I:C.

**Supplemental Table 2. Primers employed in quantitative RT-PCR.** Primers were synthesized by Sigma and used at 50μM concentration. F = forward; R = reverse.

**Supplemental Table 3. Primers employed in quantitative RT-PCR for enhancer RNA.** Enhancer regions were identified either upstream ('U') or downstream ('D') of the target gene based on presence of p300 and $H3K4me^1$, as well as presence within the enhancer-promoter unit (EPU). Primers were synthesized by Sigma and used at 50μM concentration. F = forward; R = reverse.

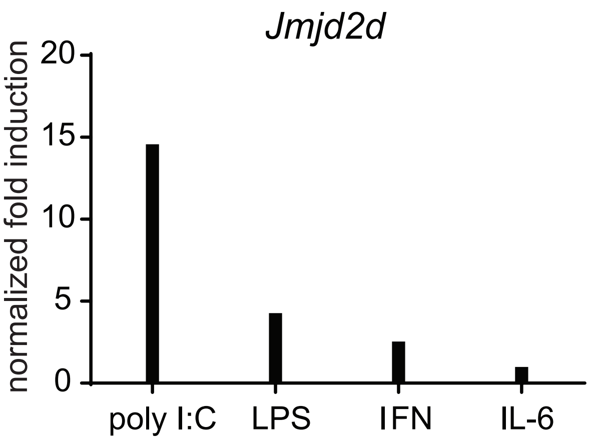


**Supplemental Figure 1**

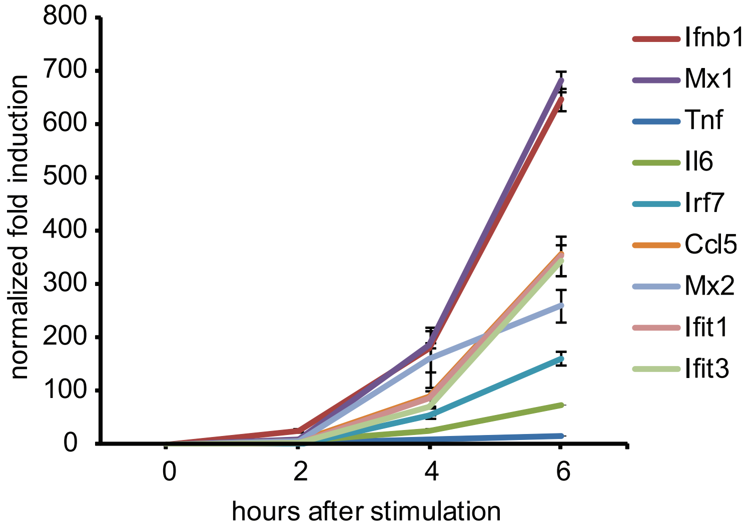


**Supplemental Figure 2**

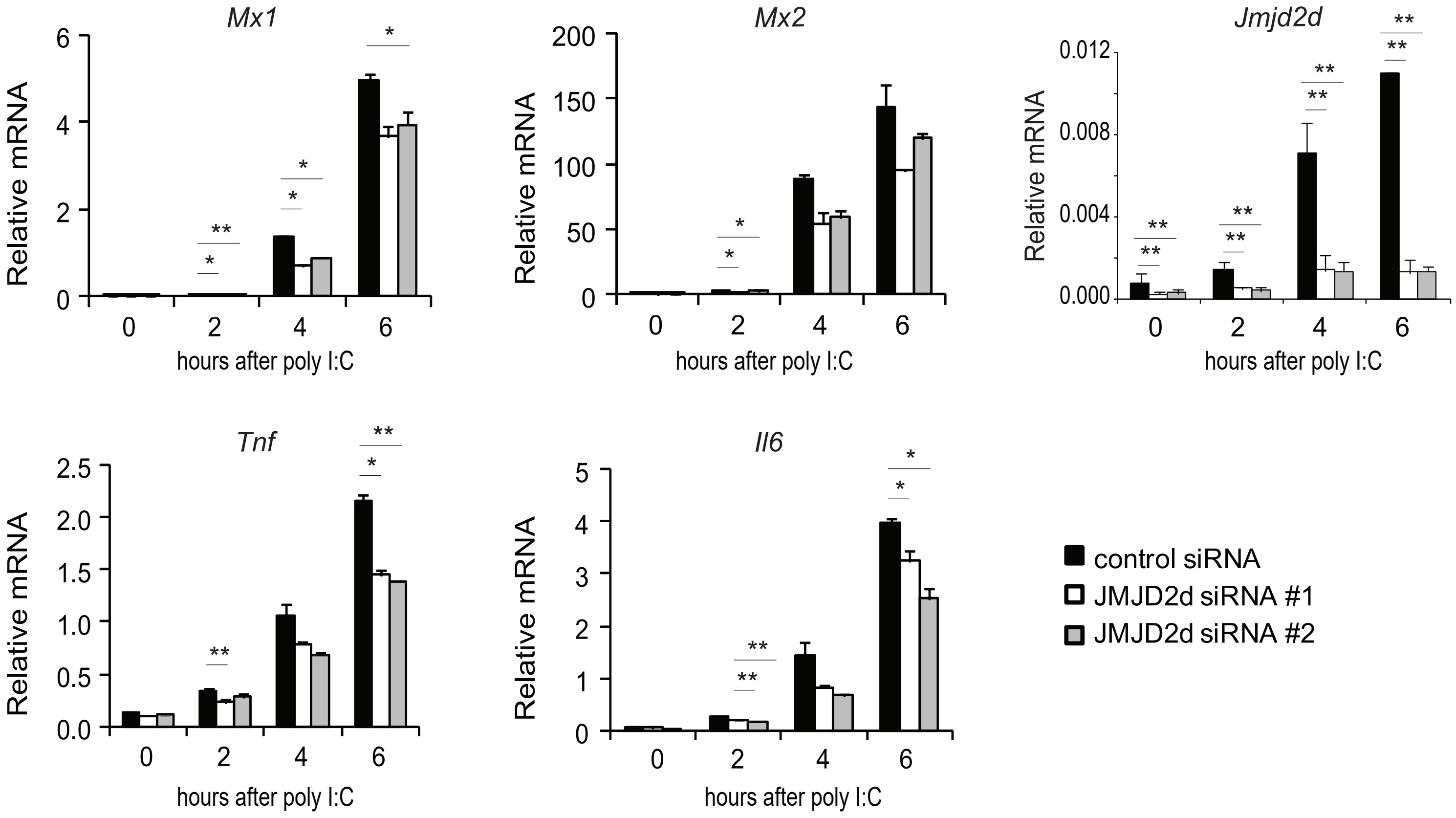


**Supplemental Figure 3**

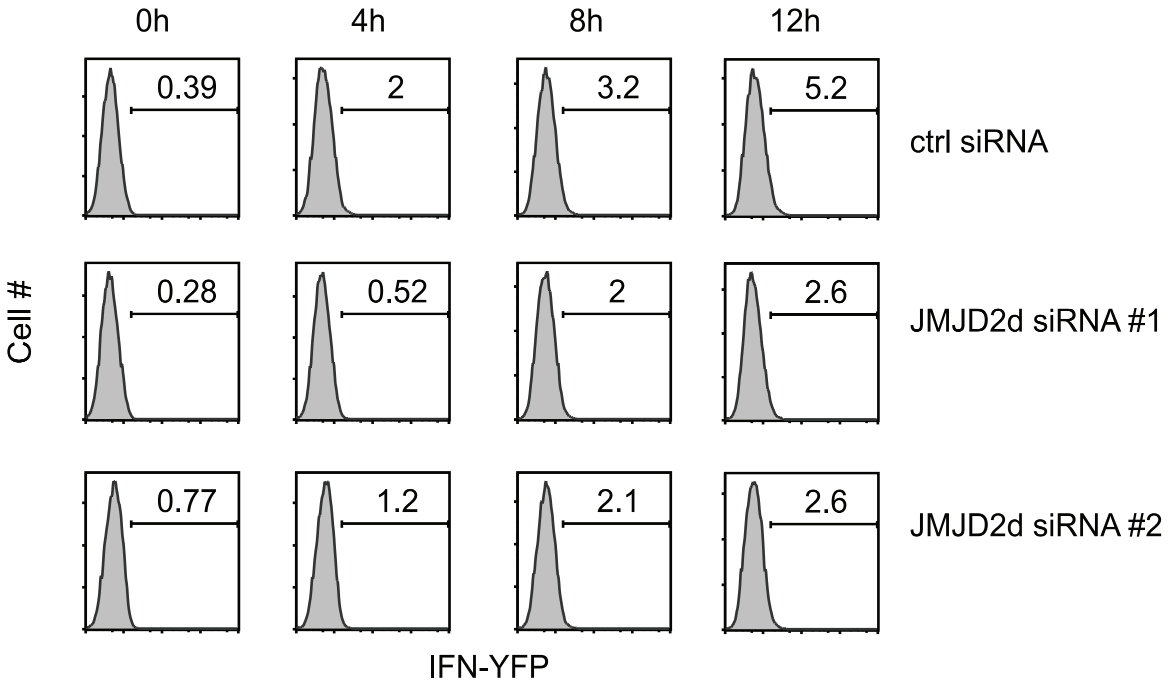


**Supplemental Figure 4**

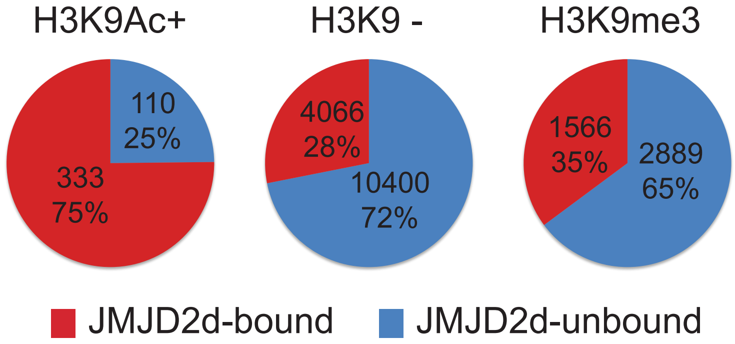


**Supplemental Figure 5**

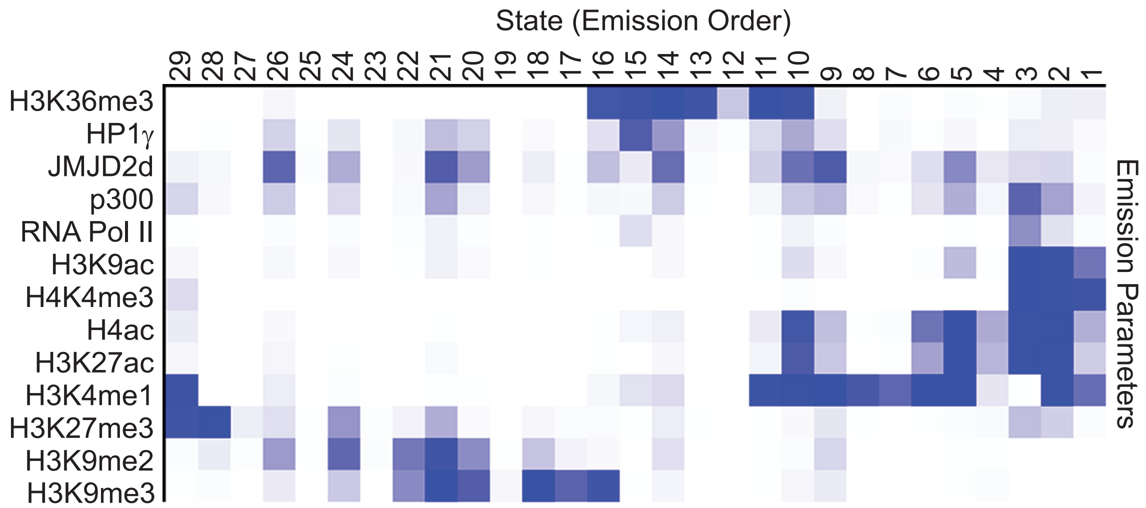


**Supplemental Figure 6**

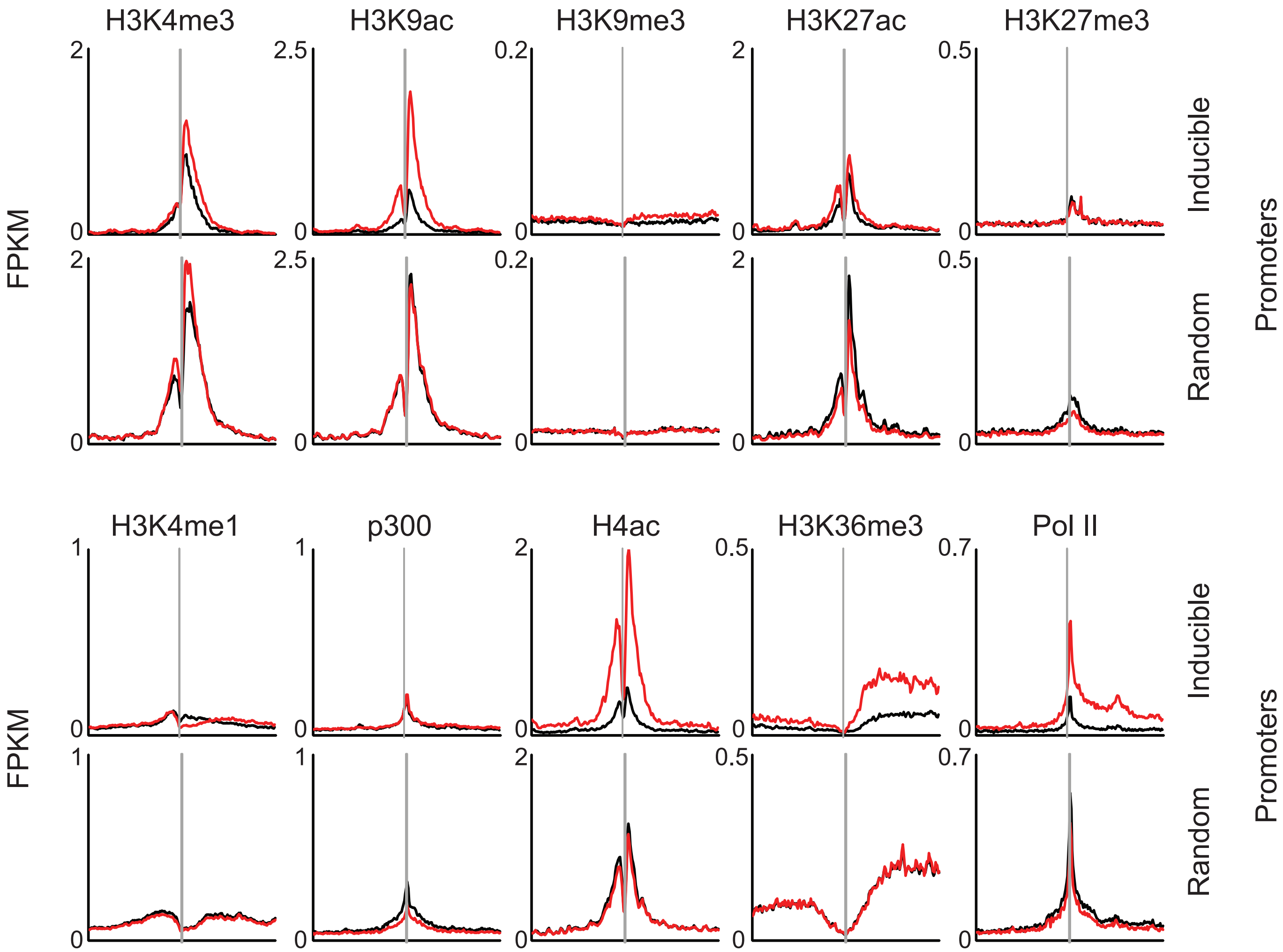


**Supplemental Figure 7**

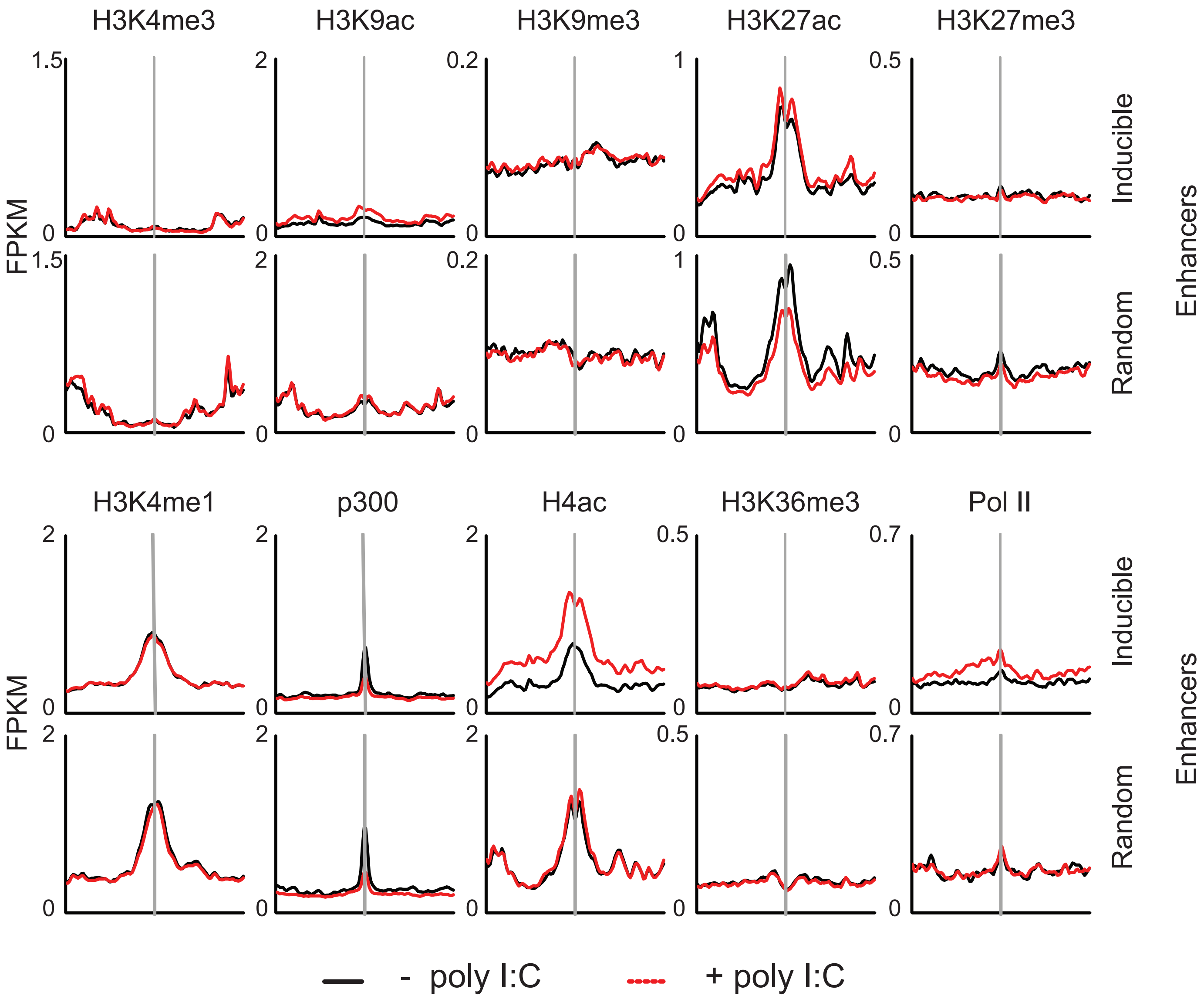


**Supplemental Figure 8**

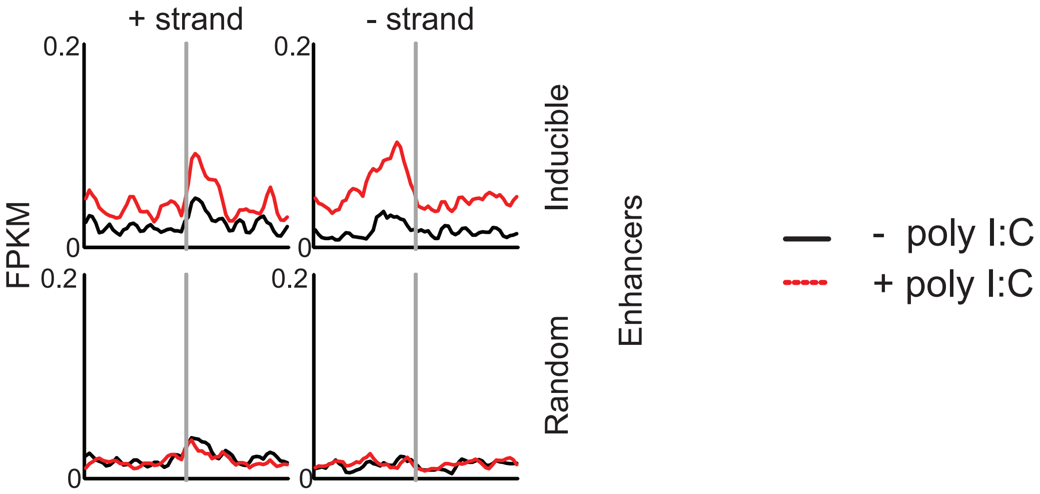


**Supplemental Figure 9**

| Gene | Fold induction | Gene | Fold induction | Gene | Fold induction | Gene | Fold induction |
|---|---|---|---|---|---|---|---|
| **Cxcl10** | 31.605846 | **Chac1** | 4.8086157 | **Hist1h1c** | 3.1748059 | **Zc3h6** | 2.4050522 |
| **Ifnb1** | 28.197994 | **Atf3** | 4.635319 | **Oas1b** | 3.1642835 | **Trex1** | 2.340291 |
| **Mx2** | 21.680067 | **Samd9l** | 4.5879884 | **Stat1** | 3.1558104 | **Relb** | 2.3122492 |
| **Ifit3** | 17.80783 | **Cd274** | 4.513661 | **Errfi1** | 3.155 | **Irf9** | 2.3026154 |
| **Ccl5** | 15.489736 | **Stat2** | 4.22059 | **Hist1h4i** | 3.0815237 | **Klf6** | 2.2975144 |
| **Usp18** | 13.926942 | **Apol9b** | 4.131248 | **Zfp36** | 3.0594335 | **Eif2ak2** | 2.2964149 |
| **Ccl2** | 10.3806505 | **Hist1h2bj** | 4.1037383 | **Slc2a6** | 3.0403197 | **Hap1** | 2.2832747 |
| **Rsad2** | 9.976725 | **Junb** | 4.0334687 | **Dusp8** | 2.954846 | **Plekha4** | 2.2785983 |
| **Gbp3** | 9.908558 | **Gbp6** | 3.9819858 | **Phlda1** | 2.920562 | **Hist1h4j** | 2.2728412 |
| **Gbp2** | 9.564882 | **Gadd45g** | 3.976631 | **Egr2** | 2.9192994 | **Ccrn4l** | 2.2654698 |
| **Tnfaip3** | 9.252282 | **Hist1h2bh** | 3.916566 | **Jun** | 2.9140236 | **Ube1l** | 2.2620642 |
| **Ifit3** | 8.9953575 | **Myd116** | 3.9051251 | **Oas1g** | 2.867266 | **Gadd45a** | 2.2534916 |
| **Iigp2** | 8.912015 | **Nfkbie** | 3.8878582 | **Dhx58** | 2.8625565 | **Ccrn4l** | 2.2346642 |
| **Cxcl1** | 8.762168 | **Gadd45b** | 3.8755217 | **Gadd45a** | 2.8510737 | **Irf7** | 2.2253335 |
| **Gbp3** | 8.508837 | **Tyki** | 3.8577104 | **Ifi47** | 2.800076 | **Daxx** | 2.2144337 |
| **Igtp** | 8.133124 | **Oasl1** | 3.6770806 | **Klf6** | 2.7642763 | **Ppm1k** | 2.1887937 |
| **Cxcl9** | 7.614855 | **Hist1h2bf** | 3.6752925 | **Hist1h4f** | 2.7262323 | **Cish** | 2.1771824 |
| **Irf1** | 6.723368 | **Map3k14** | 3.601361 | **Fos** | 2.628029 | **Apobec1** | 2.166548 |
| **Egr1** | 6.355163 | **Hist1h2bc** | 3.5996723 | **Ier3** | 2.5963612 | **Casp4** | 2.1571677 |
| **Icam1** | 6.2216988 | **Irf9** | 3.5742452 | **Ddx58** | 2.5943835 | **Ch25h** | 2.1433632 |
| **Oasl2** | 6.16569 | **Oas1b** | 3.5114527 | **Hdc** | 2.57001 | **Rhob** | 2.1350336 |
| **Ccl7** | 5.59689 | **Hist1h2bm** | 3.499247 | **Slc25a25** | 2.5684092 | **Stat1** | 2.099436 |
| **Oasl1** | 5.513505 | **Hist1h2bn** | 3.4989486 | **Ripk2** | 2.5577598 | **D14Ertd66** | 2.1451836 |
| **Axud1** | 5.478931 | **Taf15** | 3.478466 | **Ccrl2** | 2.5141284 | **Birc2** | 2.0871625 |
| **Tlr2** | 5.2830234 | **Ifit2** | 3.4701445 | **Josd3** | 2.4459813 | **Dusp6** | 2.0710497 |
| **Irgm1** | 5.151454 | **Nfkbia** | 3.3646343 | **Klf2** | 2.4446518 | **Arc** | 2.01379 |
| **Trim21** | 5.1163 | **Ifit2** | 3.3320348 | **Txnip** | 2.4422612 | | |
| **Usp18** | 5.07427 | **Hist1h2bk** | 3.3153229 | **Il15** | 2.4303243 | | |
| **Parp14** | 5.0238853 | **Clec2d** | 3.2698836 | **Hist1h2ac** | 2.4291928 | | |

### Supplemental Table 1

| | |
|---|---|
| HPRT F | CTCCTCAGACCGCTTTTTGC |
| HPRT R | TAACCTGGTTCATCATCGCTAATC |
| Cxcl1 F1 | CTTGAAGGTGTTGCCCTCAG |
| Cxcl1 F2 | GCACCCAAACCGAAGTCATA |
| Cxcl1 R | AGGTGCCATCAGAGCAGTCT |
| Cxcl2 F1 | GCCAAGGGTTGACTTCAAGA |
| Cxcl2 R1 | CTTCAGGGTCAAGGCAAACTT |
| Cxcl2 F2 | CTCCAGACTCCAGCCACACT |
| Cxcl2 R2 | AGGGTCTTCAGGCATTGACA |
| IFNb1 F | TCAGAATGAGTGGTGGTTGC |
| IFNb1 R | GACCTTTCAAATGCAGTAGATT |
| Mx1 F | GTGGTAGTCCCCAGCAATGT |
| Mx1 R | AGCACCTCTGTCCACCAGAT |
| TNF F | CCCCAAAGGGATGAGAAGTT |
| TNF R | CTCCTCCACTTGGTGGTTTG |
| IL-6 F | TGGGAAATCGTGGAAATGAG |
| IL-6 R | CCAGTTTGGTAGCATCCATCA |

### Supplemental Table 2

| | |
|---|---|
| Ifnb1 D1 F | TCAAAGAAGGGCACCACCTA |
| Ifnb1 D2 R | GTGCTGGAGGAAGGAACAACT |
| Ifnb1 D2 F | ACTGCACGCAGAGAGGTTTC |
| Ifnb1 D2 R | GGAGGTAACTGGTTGCACTGA |
| Ifnb1 U1 F | CCATCCTGTCCCTGACAGAC |
| Ifnb1 U1 R | ATGAACGAGAAAGCAGCTGTG |
| Ifnb1 U2 F | TAGAACAAACGGGGCAAAGA |
| Ifnb1 U2 R | AGATCCTGCAGTTGTGCTCAG |
| Ifnb1 D3 F | TTCAAACATTGGCCATCTGA |
| Ifnb1 D3 R | CAAGACTGAGGGTGCGTATGT |
| Mx2 U1 F | ATCAGAGCACTGGGTGTCATC |
| Mx2 U1 R | TGGTTCCTGGCATACAATGTT |
| Mx2 U2 F | ACCCTTCCACCCATCCTCTA |
| Mx2 U2 R | TCCTTGCCTCTGCAGTGTTT |
| Ccl5 U1 F | CCAACCACATTCAGACCAGAG |
| Ccl5 U1 R | TCAGAGAGCAAGTGGGTGTG |
| Ccl5 U2 F | ACCTCTGGGACAGCAAGTAGC |
| Ccl5 U2 R | AGGGCAGGCAACTAGAGACA |
| Ccl5 U3 F | CCATGGTCACAGGGTAACAAC |
| Ccl5 U3 R | CCCAGCCCTTGCTTGTATTA |
| Ccl5 U4 F | TGCCTGTCTCTGCAACAAGA |
| Ccl5 U4 R | AACCCCCTGACTCCACCTAC |
| Cxcl1 U1 F | GGCCACAGCTTCATTAAAACA |
| Cxcl1 U1 R | AAAGTGGAATCCTGGGAGACA |
| Cxcl1 U2 F | CATTCAACCTCAGTCCCATGA |
| Cxcl1 U2 R | TTGTCACAGATCCGGAGAAAG |
| Actb U1 F | ACGGAAGGGAAAGGAAAGAA |
| Actb U1 R | CCCCAGACAGTTACCACACAA |
| Actb U2 F | AAAACAAGCCAGGCACACAT |
| Actb U2 R | TAGCTGCCCTGGAACTCACT |
| Actb U3 F | TCTCTCCAGGCCCTGTTAAGT |
| Actb U3 R | CCTCTCGAGTGCTGGGATTA |

### Supplemental Table 3
